# Pico145 inhibits TRPC4-mediated mI_CAT_ and postprandial small intestinal motility

**DOI:** 10.1101/2023.08.07.552165

**Authors:** Dariia O. Dryn, Mariia I. Melnyk, Robin S. Bon, David J. Beech, Alexander V. Zholos

## Abstract

**Background & Aims:** In intestinal smooth muscle cells, receptor-operated TRPC4 are responsible for the majority of muscarinic receptor cation current (mI_CAT_), which initiates cholinergic excitation-contraction coupling. Our aim was to examine the effects of the TRPC4 inhibitor Pico145 on mI_CAT_ and Ca^2+^ signalling in mouse ileal myocytes, and on intestinal motility.

**Methods:** Ileal myocytes freshly isolated from two month-old male BALB/c mice were used for patch-clamp recordings of whole-cell currents and for intracellular Ca^2+^ imaging using Fura-2. Functional assessment of Pico145’s effects was carried out by standard *in vitro* tensiometry, *ex vivo* video recordings and *in vivo* postprandial intestinal transit measurements using carmine red.

**Results:** Carbachol (50 µM)-induced mI_CAT_ was strongly inhibited by Pico145 starting from 1 pM. The IC_50_ value for the inhibitory effect of Pico145 on this current evoked by intracellularly applied GTPγS (200 µM), and thus lacking desensitisation, was found to be 3.1 pM, while carbachol-induced intracellular Ca^2+^ rises were inhibited with IC_50_ of 2.7 pM. In contrast, the current activated by direct TRPC4 agonist (-)-englerin A was less sensitive to the action of Pico145 that caused only ∼43% current inhibition at 100 pM. The inhibitory effect developed rather slowly and it was potentiated by membrane depolarisation. In functional assays, Pico145 produced concentration-dependent suppression of both spontaneous and carbachol-evoked intestinal smooth muscle contractions and delayed postprandial intestinal transit.

**Conclusions:** Pico145 is a potent GI-active small-molecule which completely inhibits mI_CAT_ at picomolar concentrations and which is as effective as *trpc4* gene deficiency in *in vivo* intestinal motility tests.

## Introduction

Acetylcholine is the primary excitatory neurotransmitter in the gut. When released by enteric motor neurons, it controls excitation-contraction coupling and complex patterns of motility of the gastrointestinal (GI) tract by acting on muscarinic receptors expressed both in interstitial cells of Cajal and smooth muscle (SM) cells^1, 2^. In SM cells, M2 and M3 muscarinic receptors are present, predominantly coupled to G_i/o_ and G_q/11_ proteins, respectively. Cholinergic GI SM contraction is predominantly an M3 response mediated by the G_q/11_/phospholipase C (PLC) pathway causing breakdown of phosphatidylinositol 4,5-bisphosphate (PIP_2_) into 1,2-diacyl-sn-glycerol and D-myo-inositol 1,4,5-trisphosphate (InsP_3_). InsP_3_ directly initiates myocyte contraction by rapidly releasing Ca^2+^ from the sarcoplasmic reticulum via InsP_3_ receptors. The cholinergic contraction develops in parallel with membrane depolarisation, the generation of slow waves and accelerated action potential discharge^1, 3^. Thus, concomitant activation of L-type Ca^2+^ channels is required for the cholinergic contraction to develop, while muscarinic agonist-evoked Ca^2+^ release may contribute to the contraction indirectly via potentiation of the electrical membrane responses^4^. Indeed, activation of muscarinic receptors causes multiple effects on the activity of many types of ion channels^1, 5–7^. Among these, the muscarinic cation current (mI_CAT_) mediated mainly by TRPC4 (∼85%) and, to a much lesser extent, TRPC6 channels is of particular interest. mI_CAT_ is the principal component of cholinergic excitation-contraction coupling^8^.

TRPC4 channels are activated synergistically by both M2 and M3 muscarinic receptors resulting in smooth muscle depolarization, activation of L-type Ca^2+^ channels and contractile response. Importantly, the roles of M2 and M3 receptors in mI_CAT_ activation are not simply additive. Although mI_CAT_ behavior was most consistent with the pharmacological profile of M2 receptors, activation of M3 receptors was also required in a permissive manner^9–11^. The M2 signalling arm acts via a presumably direct binding of activated Gα_i/o_ subunits (most prominently Gα_i2_) to the intracellular domain of the channel protein^12, 13^. The M3/PLC/G_q/11_ arm of the cellular signalling leading to mI_CAT_ activation is not completely understood but tonic inhibition of TRPC4 by PIP_2_^14^ may explain the prerequisite for the reduction of PIP_2_ levels for efficient channel opening at least in case of the full-length TRPC4α isoform^15^. Moreover, mI_CAT_ is strongly potentiated by intracellular Ca^2+^ release, with InsP_3_ receptors playing a central role in this process^16^.

Therapeutically, both M3-selective and non-selective muscarinic antagonists are traditionally studied and clinically used for treating disorders of smooth muscle hyperactivity^17^. Since anticholinergics cause multiple side effects, such as dry mouth and blurred vision due to M3 inhibition or tachycardia resulting from M2 blockade, inhibition of mI_CAT_ as a focal point may be a therapeutically beneficial alternative. Further highlighting the possible clinical benefits of mI_CAT_ downregulation, physiologically mI_CAT_ is kept in check by several endogenous factors, including current desensitisation, which is dependent on protein kinase C activation^18^, PIP_2_ levels^15^, and polyamines^19^. If some of these multiple endogenous regulatory mechanisms fail under pathophysiological conditions, a pharmacological intervention aimed at TRPC4 may indeed be beneficial.

Recent progress with the development of TRPC1/4/5 pharmacology has provided new chemical probes for detailed studies of TRPC1/4/5 channels (for critical reviews, see^20, 21^), as well as the clinical candidates BI-1358894 and GFB-887 (currently undergoing Phase IIa trials for the treatment of CNS disorders by Boehringer Ingelheim and Karuna Therapeutics, respectively)^22, 23^. The most promising compounds include the xanthine derivatives Pico145 (HC-608) and HC-070, which have been thoroughly characterised in terms of effects on different TRPC1/4/5 tetramers, selectivity, and suitability for use in cells, tissues and animals^20, 24, 25^. Here, we report our studies of Pico145’s effect on TRPC4 channels coupled to muscarinic receptors in mouse ileal cells, as well as assessment of intestinal function *in vitro*, *ex vivo* and *in vivo*.

## Results

### Inhibitory effects of Pico145 on carbachol- and GTPγS-activated current

In our first electrophysiology experiments, carbachol was applied at sub-maximal concentration of 50 µM^9^, while 1 mM GTP was added to the pipette solution to minimize current desensitization. Mean mI_CAT_ amplitude at −40 mV was −620.6±42.5 pA (n=34). In all cells tested, Pico145 strongly inhibited the current, starting from concentrations as low as 1 pM (Figure 1). Our voltage protocol (Figure 1A, top) allowed us to monitor steady-state current amplitudes at three test potentials (−40, −120 and 80 mV) for time-course plots (Figure 1B,D,F,H), kinetics of current deactivation at −120 mV and reactivation upon stepping back to −40 mV, as well as steady-state I-V relationships measured by slow, 6 s duration voltage ramps (Figure 1C,E,G).

**Figure 1.**
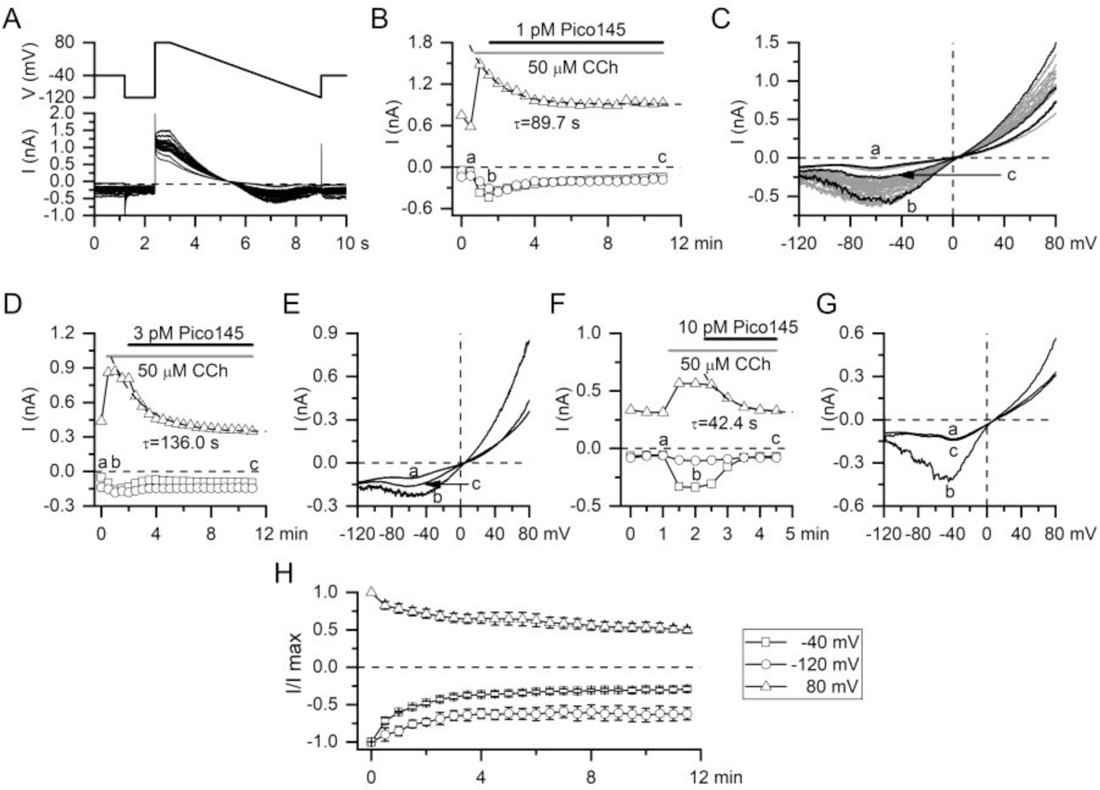
Pico145 strongly inhibits carbachol-induced mI_CAT_ in isolated mouse ileal myocytes. (*A*) Top, voltage protocol used for mI_CAT_ recordings; bottom, superimposed current traces recorded in control, after 50 μM carbachol application and following Pico145 application at 1 pM. (*B*) Time course of mI_CAT_ amplitudes measured at the end of voltage steps (by averaging the last 100 ms segment of the trace) to −40, −120 and 80 mV, as indicated (symbols as in panel H). Pico145-induced inhibition of mI_CAT_ progressed exponentially with time constants of 89.7 ms at 80 mV as shown by the fitted dotted line. (*C*) Steady-state I-V relationships of mI_CAT_ measured in control (a), at the peak response to 50 μM carbachol (b) and at the steady-state inhibition by Pico145 (c). These moments are indicated in panel B. Panels D-F and G-H illustrate similar experiments, in which Pico145 was applied at 3 and 10 pM, respectively. *(H)* Mean normalised data showing time course of mI_CAT_ amplitudes in control (n=3).

Pico145-induced inhibition of mI_CAT_ developed rather slowly. The process could be fitted by a single exponential function with time constants in the range of tens of seconds (superimposed dotted lines in Figure 1B,D,F). Pico145 applied at 10 pM almost abolished mI_CAT_ (Figure 1F,G). Pico145 wash-out was not feasible because the patch-clamp gigaseal became unstable during long-duration recordings that were needed to observe its effect.

Ongoing receptor desensitization resulted in substantially reduced mI_CAT_ during similar period of time (Figure 1H), preventing accurate quantification of the Pico145 concentration-response relation for carbachol-induced mI_CAT_. Thus, to avoid current desensitisation, we used the stable GTP analogue GTPγS to directly activate mI_CAT_. GTPγS (200 μM) was infused into the cells via the patch-pipette. This resulted in slow mI_CAT_ development, which peaked in about 3-5 min after membrane break-through (Figure 2A,B).

**Figure 2.**
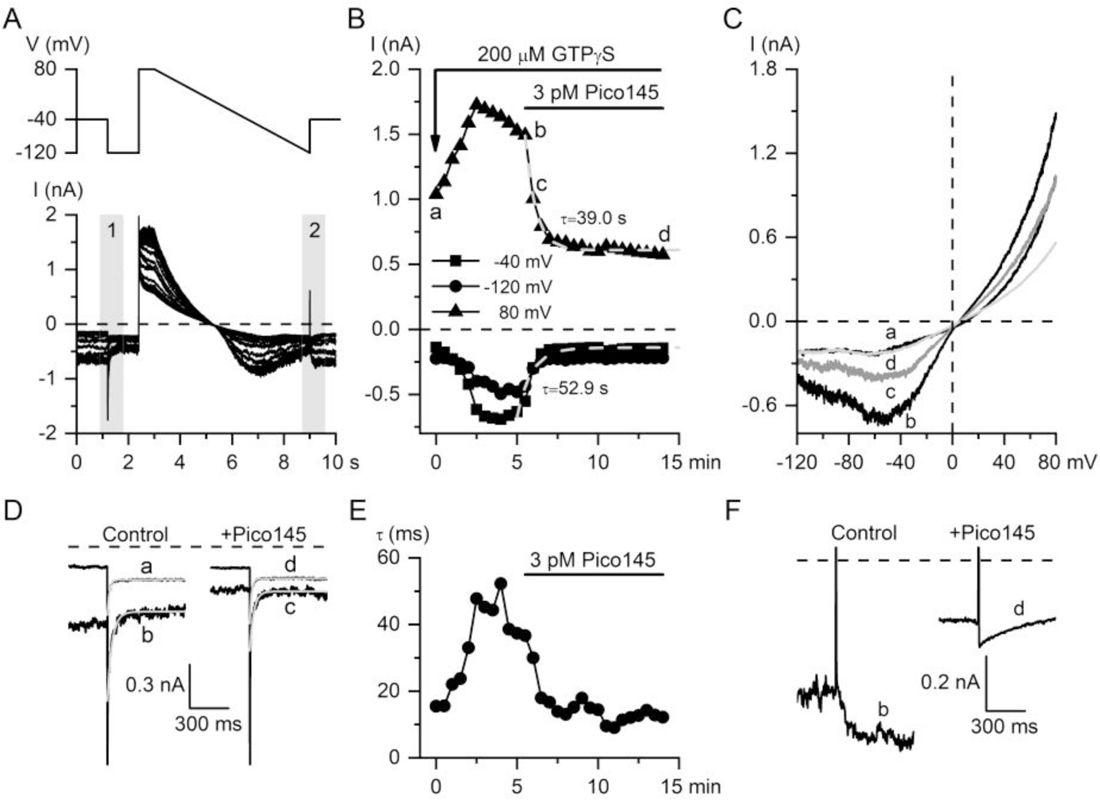
Pico145 strongly inhibits mI_CAT_ induced by GTPγS, thus bypassing activation of muscarinic receptors. (*A*) Top, voltage protocol used for mI_CAT_ recordings; bottom, corresponding superimposed current traces recorded starting shortly after break-through with patch pipette containing 200 μM GTPγS. Pico145 at 3 pM was applied after mI_CAT_ reached its maximal amplitude. (*B*) Time course of mI_CAT_ amplitudes at three different test potentials, as indicated. Pico145-induced inhibition of mI_CAT_ developed exponentially with time constants of 52.9 and 39.0 s at −40 and 80 mV, respectively, as shown by the fitted dotted lines. (*C*) Steady-state I-V relationships of mI_CAT_ measured immediately after break-through (a - control), at the peak response to GTPγS (b) and in the presence of Pico145 for partially (c) and fully inhibited (d) currents. Corresponding moments are indicated in panel B. *(D)* Voltage-dependent current deactivation during voltage steps from −40 to −120 mV (shaded area 1 in panel A) during current development (left panel) and during its inhibition by Pico145 (right panel). Smooth light gray lines show single exponential fit of the deactivation kinetics. *(E)* Corresponding time constants of current decline, which represent channel mean open dwell time at −40 mV. *(F)* Voltage-dependent current reactivation during voltage step from −120 to −40 mV (shaded area 2 in panel A) during current development (left panel) and during its inhibition by Pico145 (right panel).

In these experiments, mean GTPγS-induced current amplitude at −40 mV was −521.8±43.4 pA (n=30, combined statistics for all cells with GTPγS, to which different Pico145 concentrations were applied; P=0.11 compared to carbachol-induced mI_CAT_). When the activation reached a peak, Pico145 was applied at different concentrations. A typical example is shown in Figure 2. Pico145 (3 pM), inhibited the GTPγS-induced current at two different test potentials with time constants of 52.9 and 39.0 s at −40 and 80 mV, respectively (Figure 2B). Similarly to carbachol-induced currents, the I-V relations in Figure 2C show high efficacy of Pico145 in mI_CAT_ inhibition.

There are also changes in mI_CAT_ voltage-dependent kinetics during Pico145 action. Current deactivation kinetics during voltage step to −120 mV normally slow down with the increasing activation of G-proteins. This indicates a prolongation of mean open dwell time of the channel^26^. However, deactivation kinetics were accelerated in the presence of Pico145 (Figure 2D,E). Mean deactivation time constant was 34.0±4.1 ms in control, decreasing to 13.7±2.7 ms after 3 pM Pico145 application (n=6, P<0.01 by paired *t*-test). Moreover, stepping to −40 mV at the end of the voltage ramp, i.e. from −120 mV, normally causes voltage-dependent activation of mI_CAT_, but in the presence of the inhibitor these current relaxations changed their direction (Figure 2F), indicating additional voltage-dependent current inhibition.

For quantitative pharmacological assessment, maximal GTPγS-induced inward current at the holding potential of −40 mV was normalised as 1.0 in each cell. Mean data fitted by Hill-type equation in Figure 3B revealed an IC_50_ value of 3.1±0.2 pM and a Hill slope of 0.70±0.03 (n ≥ 5 for each concentration in the range of 0.1-100 pM). Our results show that Pico145 is a potent and efficacious inhibitor of mI_CAT_, independent of the involvement of muscarinic receptors.

**Figure 3.**
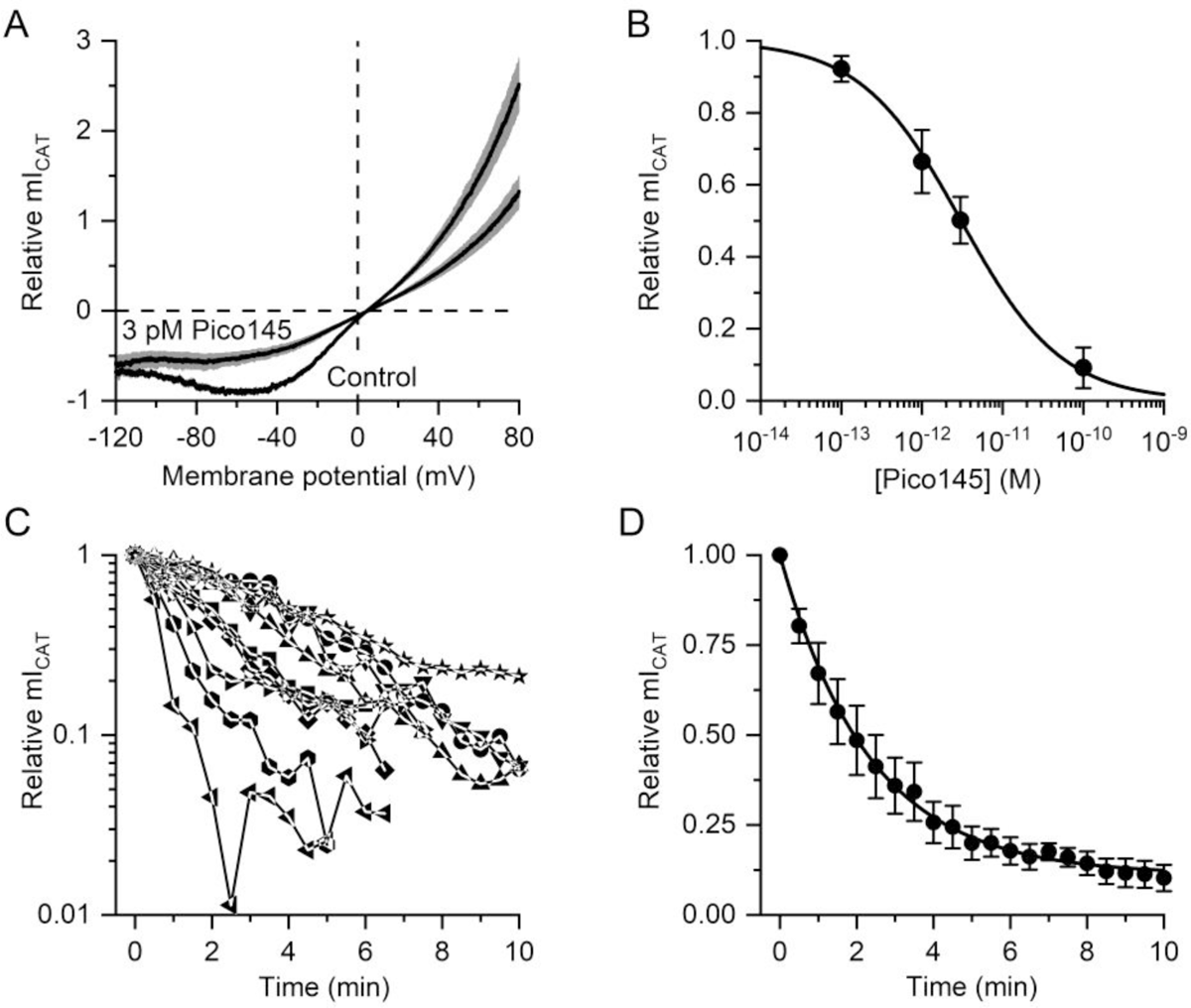
Concentration and time dependence of the inhibitory effect of Pico145 on GTPγS-induced mI_CAT_. (*A*) Mean I-V relationships of mI_CAT_ measured at its full activation by intracellular 200 μM GTPγS application in control and at its steady-state inhibition by 3 pM Pico145. In each cell, maximal inward current was normalised as 1.0. This corresponded to −652.9±96.9 pA (n=5). The grey band shows the SEM values. (*B*) Dose-response data points for the inhibitory action of Pico145 on GTPγS-induced mI_CAT_ measured at −40 mV were fitted by the Hill equation with the best-fit values: IC_50_=3.1±0.2 pM, Hill slope of 0.70±0.03. (*C*) In each individual ileal myocyte, as denoted by different symbols, amplitude of mI_CAT_ induced by intracellular 200 μM GTPγS application at 80 mV before 100 pM Pico145 application at time 0 was normalised as 1.0 (this corresponded to 1197. 9±161.0 pA on average, n=9) and plotted on a semi-logarithmic scale. (*D*) Mean data were fitted with a single exponential function with the time constant of 140±6 s.

Slow inhibition of the current was at odds with high potency of the drug. We therefore examined this phenomenon in more detail. Even at 100 pM, Pico145 inhibited mI_CAT_ recorded at −40 mV slowly. In individual cells (represented by different symbols in Figure 3C), the inhibition time constant ranged from 37.5 to 666.7 s (n=9). When mean normalised values of the current were fitted, the mean time constant was 140.0±5.7 s (Figure 3D, n=9).

### Voltage-dependence of mI_CAT_ inhibition by Pico145

Voltage dependence of the effect of Pico145 on TRPC4-mediated mI_CAT_ is in agreement with previous findings in TRPC4-expressing HEK T-REx cells^24^. This implies that Pico145 binds within the membrane potential field, consistent with our cryoEM data for the TRPC5:Pico145 complex^27^). To further characterise the effects of Pico145, we investigated alterations in voltage-dependence of current kinetics. During voltage step from a very negative potential to very positive potentials an additional current activation is normally seen (Figure 4A). In the presence of Pico145, additional current inhibition developed, seen as negatively going current relaxation. The same behaviour was also observed for Pico145 inhibition of GTPγS-induced currents (Figure 4B). In the latter case, the mean time constant of current decline was 107.4±11.1 ms (n=13) and the kinetics were independent of Pico145 concentration: the time constants were 99.8±8.4 ms (n=4) at 1 pM *vs.* 112.9±25.3 ms (n=5) at 10 pM of Pico145 (P=0.67). However, if voltage steps to 80 mV were applied from a holding potential of −40 mV, such current relaxations at 80 mV were no longer present (Figure 4C).

**Figure 4.**
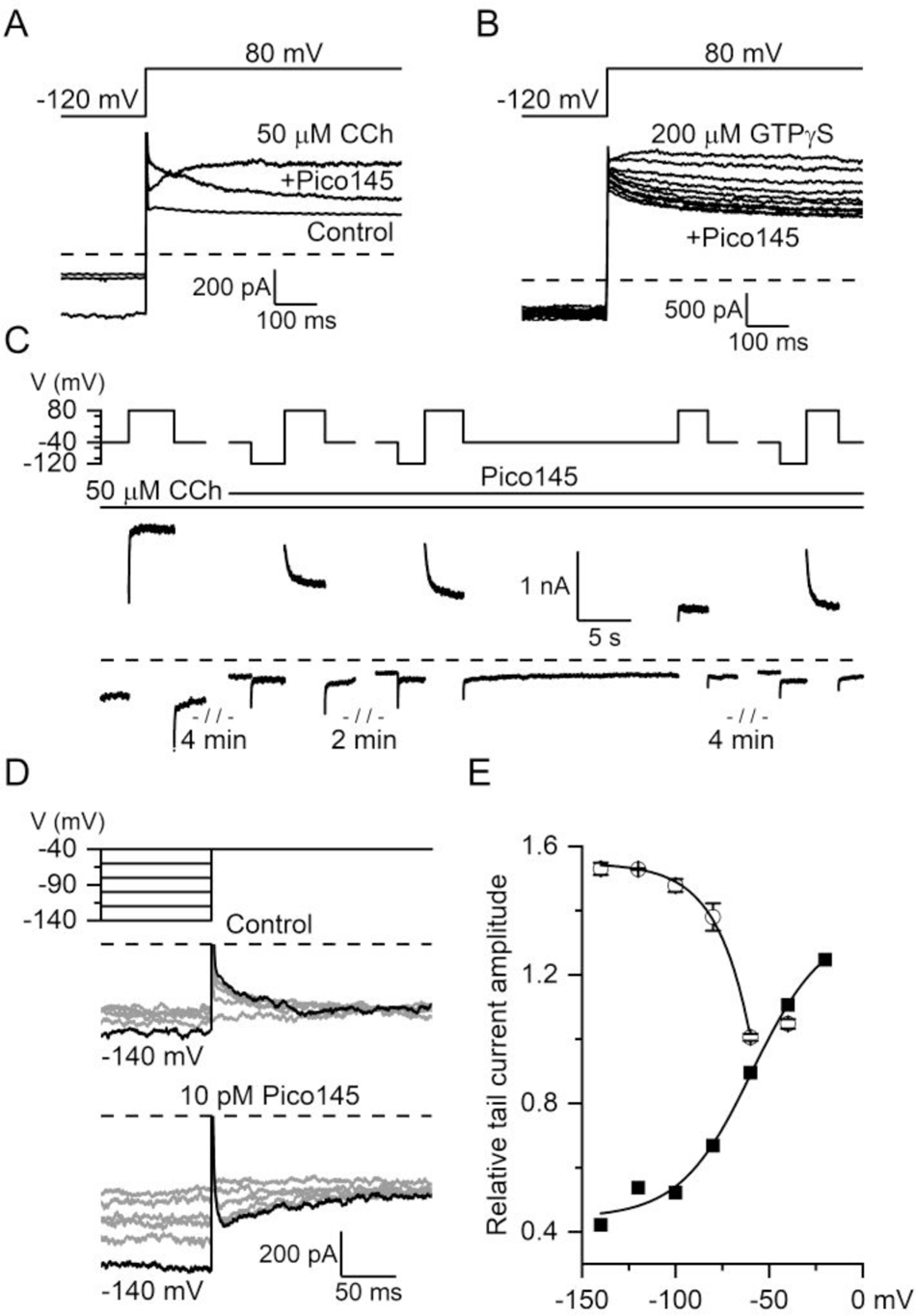
Pico145 blockage of TRPC4 is voltage-dependent. (*A & B*) Pico145 inhibition of mI_CAT_ is partially removed by voltage steps from −40 mV to −120 mV, while the subsequent step to 80 mV causes additional current inhibition seen as current relaxation not present in control, for both carbachol-(*A*) and GTPγS-induced (*B*) currents. In panel B, currents recorded at 30 s intervals after 3 pM Pico145 application are superimposed to further illustrate kinetics of the development of this effect along with progression of current inhibition. (*C*) Voltage-dependent current relaxations upon voltage steps from −120 to 80 mV become more pronounced with time of Pico145 action and these are not seen without the prior voltage step to −120 mV (compare the last 2 traces). *(D)* Double-pulse voltage step protocol (top) reveals additional mI_CAT_ inhibition by Pico145 at the holding potential of −40 mV (bottom traces), while in control voltage-dependent current activation develops after a negative step (middle traces). *(E)* Relative tail current amplitude in control (black squares) and in the presence of Pico145 (open circles). See text for more details.

It thus appeared that voltage dependence of Pico145 interaction with the channel encompassed only a certain range of potentials negative to −40 mV. To examine this question in more detail, we employed another protocol that included voltage steps from −40 to −140 mV with a −20 mV increment (Figure 4D). In control experiments, stepping from negative values to −40 mV produced inwardly going tail currents, in line with usual mI_CAT_ voltage dependence (Figure 4E; closed squares). In contrast, in the presence of Pico145, these tail currents reversed their direction (Figure 4D, bottom panel), showing additional current inhibition by Pico145 after it was disinhibited during the negative step (compare also to Figure 2F). This effect required membrane hyperpolarisation negative to about −60 mV and it developed with an e-fold increase per 17.9±0.2 mV (n=3) until reaching saturation at about −120 mV (Figure 4E, open circles).

### (-)-Englerin A-induced current is less sensitive to Pico145

Pico145 has been characterised as an inhibitor of TRPC1/4/5 channels overexpressed in HEK cells and activated by their direct agonist (-)-englerin A (EA). It was shown that Pico145 potency inversely correlated with EA concentration, consistent with Pico145 being a competitive antagonist of EA^24^. Thus, we further characterised the effect of Pico145 on EA-activated TRPC4 channels. Remarkably, there was no desensitization of EA-induced currents; instead their amplitudes even slightly increased with time (Figure 5A,B).

**Figure 5.**
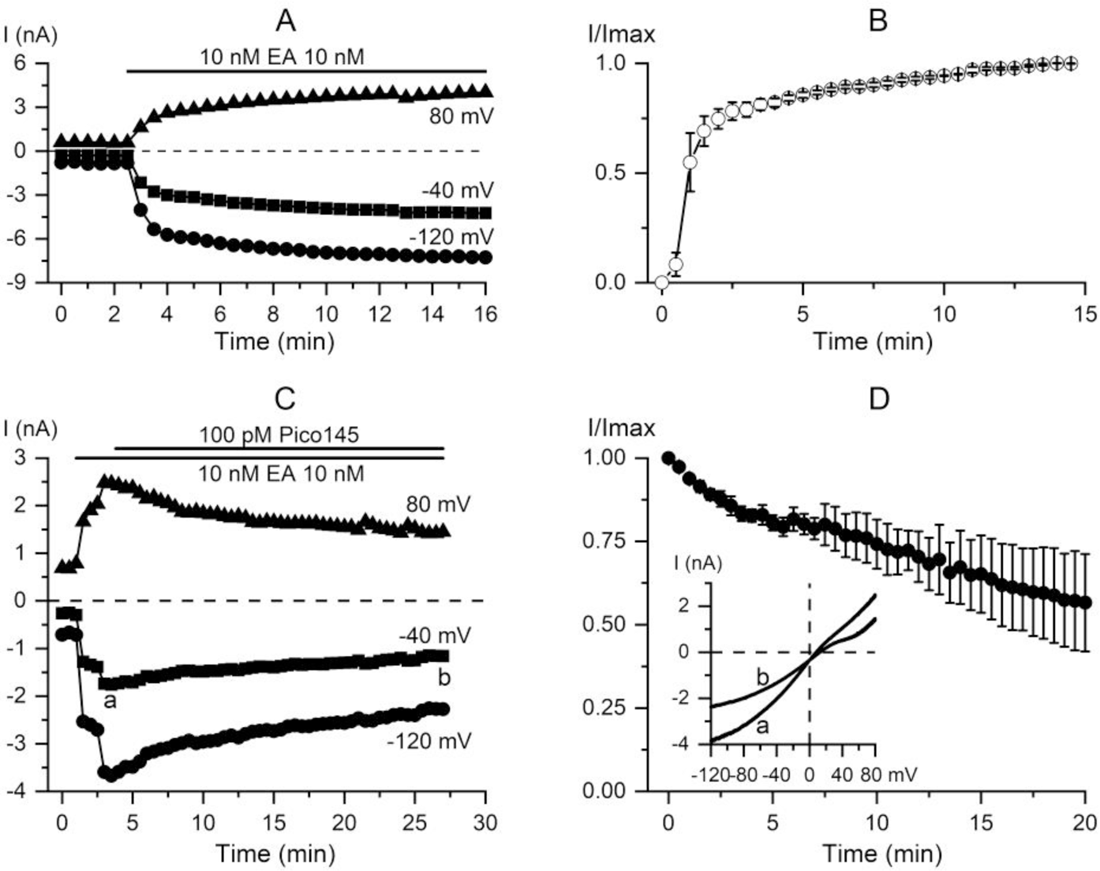
(−)-Englerin A (EA) induced currents are less sensitive to the inhibitory action of Pico145 than carbachol- or GTPγS-induced currents. (*A*) A representative example of TRPC4 activation by its direct agonist EA applied at 10 nM in control. *(B)* Mean time course of EA-induced current recorded at −40 mV. In each cell Imax was normalised as 1.0; this corresponded to −2765.0±786.8 pA on average (n=3). *(C)* Partial inhibition of EA-induced current by 100 pM Pico145. Steady-state current amplitudes were measured at the end of voltage steps to −40, −120 and 80 mV, as indicated. (*D*) Mean time course of the inhibitory effect of 100 pM Pico145 on EA-induced (10 nM) currents recorded at −40 mV (n=5). The inset shows I-V relations measured at the corresponding time points denoted in panel C.

Application of 100 pM Pico145 at the peak response to 10 nM EA caused mI_CAT_ inhibition only by 43% (Figure 5C). Maximal EA-induced current amplitude at −40 mV was −1652.6±453.2 pA (n=5), which was about 3 fold larger compared to carbachol- or GTPγS-induced currents. Amplitude of the normalised steady-state EA current recorded at −40 mV was 0.57±0.15 after its inhibition by 100 pM Pico145 (Figure 5C n=5, P<0.05). Again, the inhibition progressed slowly, with a time constant of 172.6±16.2 s, which was similar to carbachol- or GTPγS-induced currents (compare Figures 5D and 3D).

### Inhibition by Pico145 of carbachol-induced intracellular Ca^2+^ and contractile responses

In GI SM, a rise in [Ca^2+^]_i_, depending on agonist concentration and time of its action, can be either in the form of periodic oscillations, the so-called Ca^2+^ waves, or more sustained responses following an initial large transient due to InsP_3_-induced Ca^2+^ release^1^. As illustrated in Figure 6A,B in our Ca^2+^ imaging experiments using Fura-2, both types of responses were seen in cells stimulated by carbachol (n=11).

**Figure 6.**
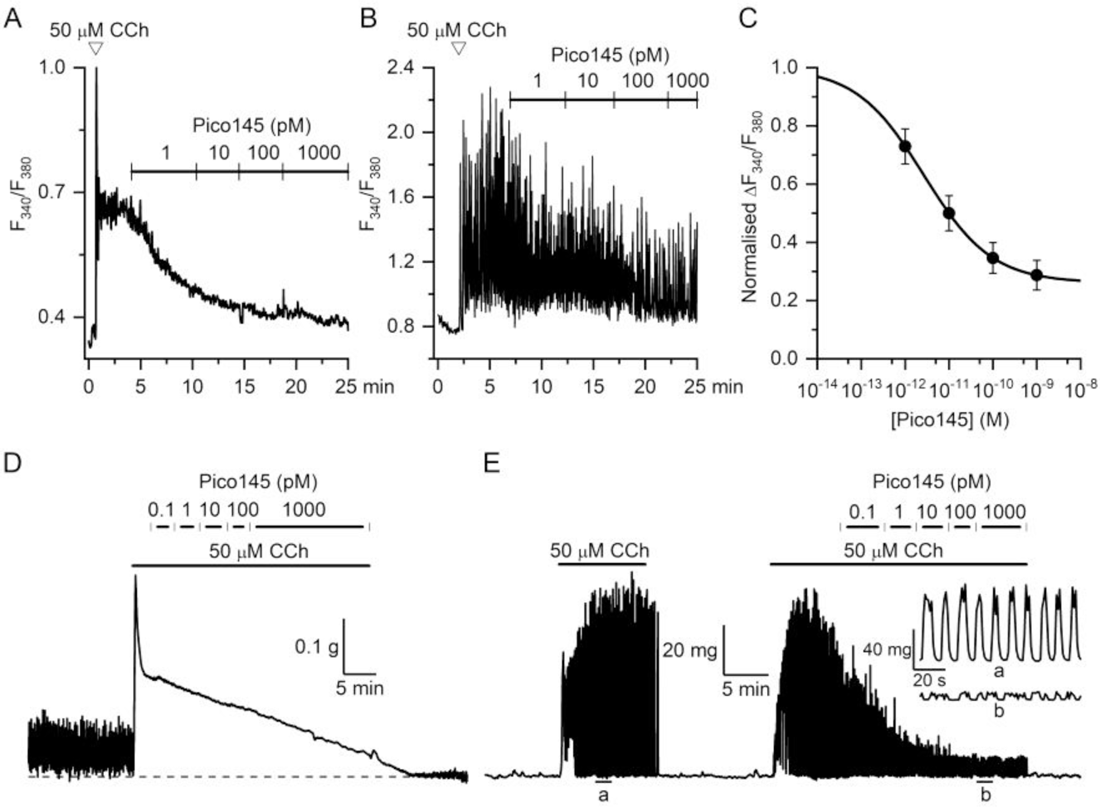
Functional assessment of Pico145 blockade of mI_CAT_ using intracellular Ca^2+^ imaging and *in vitro* tensiometry. *(A & B)* Representative examples of [Ca^2+^]_i_ responses to 50 µM carbachol applications in longitudinal ileal myocytes loaded with Fura-2 AM (n=11) and their suppression by cumulative applications of Pico145 at concentrations ranging from 1 pM to 1 nM, as indicated. (C) Dose-response data points for the inhibitory action of Pico145 on carbachol-induced [Ca^2+^]_i_ responses. Data are given as mean normalised F_340_/_380_ ratio calculated after subtracting the baseline F_340_/_380_ ratio (0.32±0.01) and normalised by the F_340_/_380_ ratio measured before Pico145 application (0.43±0.04, n=8). Data points were fitted by the logistic function with maximum constrained at 1.0 with the best-fit values: IC_50_=2.67±0.02 pM, Hill slope of 0.561±0.003. *(D & E)* Similarly, Pico145 applied at ascending concentrations ranging from 0.1 pM to 1 nM suppressed both sustained tonic component of 50 µM carbachol-induced contraction (D, longitudinal ileum), as well as transient phasic contractions (E, circular layer of the ileum).

Cumulative application of Pico145 (1-1000 pM) inhibited these carbachol-induced [Ca^2+^]_i_ responses with an IC_50_ of 2.67±0.02 pM (Hill slope 0.561±0.003; n=8; Figure 6C) Qualitatively, Pico145 was more potent in inhibiting sustained type responses (Figure 6A) compared to oscillatory type responses (Figure 6B). This is as expected since [Ca^2+^]_i_ oscillations in these cells arise mainly due to InsP_3_ signalling, while the main functional role of TRPC4 is to produce membrane depolarisation that activates a sustained Ca^2+^ entry via L-type Ca^2+^ channels.

In tensiometric recordings, both longitudinal and circular intestinal SM layers were tested. In longitudinal SM, application of 50 μM carbachol usually initiated transient phasic contraction followed by a sustained tonic contraction (Figure 6D), whereas in circular SM, periodic phasic contractions were observed (Figure 6E). Such patterns of contractile activity are consistent with the different patterns of carbachol-induced [Ca^2+^]_i_ responses (Figure 6A,B). Application of Pico145 (0.1-1000 pM) suppressed these responses. Thus, in the presence of 3 nM Pico145 the sustained tonic component of carbachol-induce contraction was reduced to 40.8±2.7% of its initial level (n=6). The remaining small amplitude phasic contractions likely occur due to oscillatory InsP_3_-induced Ca^2+^ releases, which are no longer amplified by the TRPC4/L-type Ca^2+^ channels system (Figure 6E). These results show that Pico145 suppresses [Ca^2+^]_i_ and contractile responses in GI SM. We therefore next tested if there are *in situ* and *in vivo* effects.

### Pico145 suppresses motility of ileal segments

The inhibitory action of Pico145 was clearly visualized in *ex vivo* experiments on intact ileal segments of the small intestine. In our initial tests, administration of 100 pM or 1 nM Pico145 did not affect intestinal motility, likely because of more substantial diffusion barriers present in these preparations. However, intestinal motility, both spontaneous and carbachol-induced, was suppressed by 3 nM Pico145 (Figure 7, Supplementary Figure 1, Supplementary Video 1). The movements of the ileum were represented as the MTrackJ D2S parameter, indicating the distance (in pixels) from the start point of the track to the end point (Supplementary Figure 1). Phasic contractile activity typical for small intestinal motility (Figure 7A) was inhibited by 3 nM Pico145 (Figure 7B). The mean distance of the track from 6 selected points in ileal preparation was 51.04±8.21 pixels in control, while 5 min after Pico145 application it was reduced to 28.09±2.25 pixels (P<0.002, Figure 7C). Pico145 (3 nM) similarly efficiently inhibited ileal contractions initiated by 50 µM carbachol. Figure 7D shows considerably less distance of the tracks in a preparation pre-treated with Pico145 for 5 min. The edges of the preparation were shifted about three fold less after Pico145 application.

**Figure 7.**
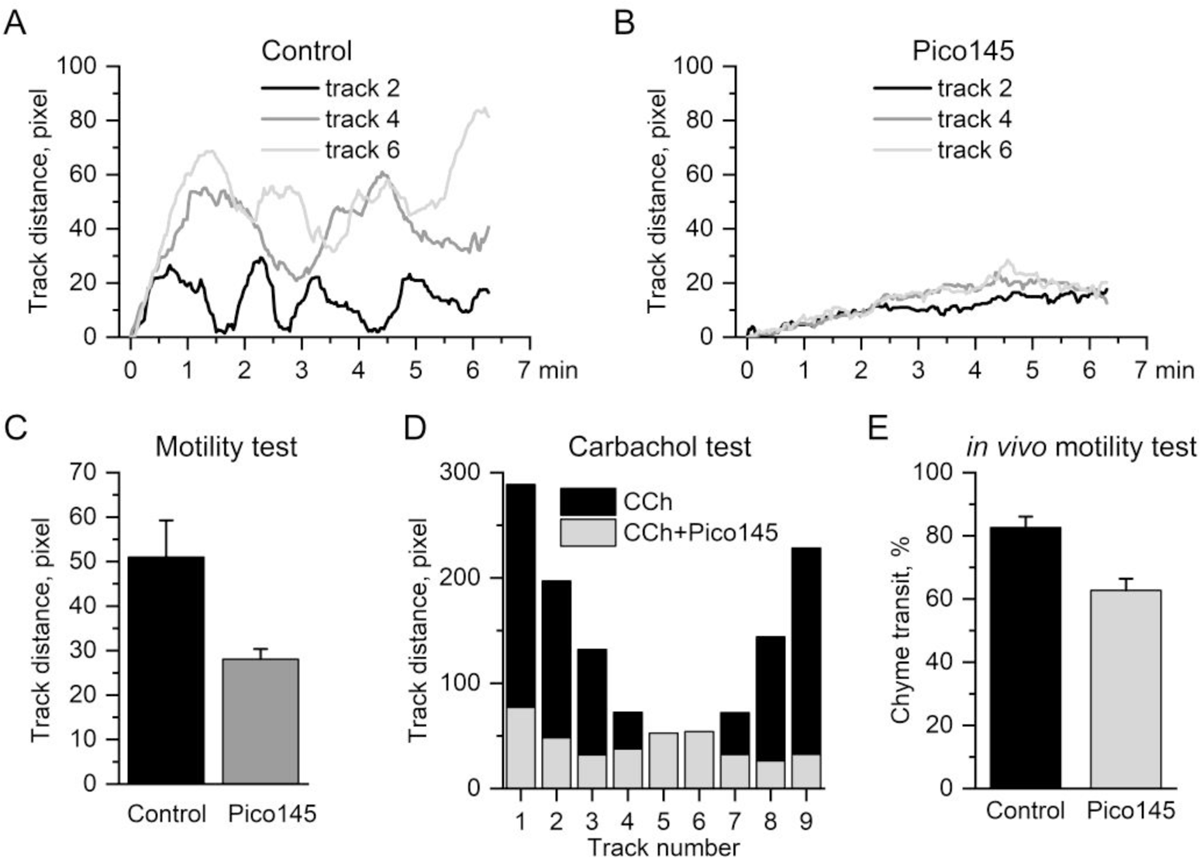
Pico145 suppresses both spontaneous and 50 µM carbachol-induced ileal contractile activity *ex vivo*, as well as postprandial small intestinal motility *in vivo*. *(A)* Spontaneous movements of intact segments of the small intestine were quantified as displacement of several selected points along the preparation (denoted tracks 2, 4 & 6 in the Supplementary Figure 1). *(B)* Pico145 (3 nM, 5 min) inhibited the contractile activity in the same preparation. *(C)* Mean track distance from the 6 selected points in control and 5 min after 3 nM Pico145 application. *(D)* Pico145 (3 nM, 5 min) inhibited carbachol-induced contractile activity represented as track distance for the 9 selected points. Note that the central part of the ileal segment (tracks 4-7) did not show much movement, while most movements inhibited by Pico145 were observed at both ends of the segment, that is at tracks denoted 1-3 and 8-9. All data are calculated from representative *ex vivo* visual experiments using the ImageJ and MTrackJ software. (*E)* Mean data illustrating Pico145-induced inhibition of small intestinal transit *in vivo*. Chyme transit was expressed as a percentage of the small intestine total length of each individual mouse in control (n=10) and after Pico145 administration at the dose of 1 mg per kg body weight.

### Small intestinal transit inhibition

To determine to what extent Pico145 could reduce small intestinal transit *in vivo*, oral gavage administration was used. Mice were fasted for 24 hours beforehand and then given either Pico145 (1 mg/kg of body weight) or 0.25 mL of distilled water (control). Then, 1 g of food (grains) was stained with carmine red (6% w/v) and both groups were fed 10 minutes after the administration of Pico145. 30 minutes later the mice were sacrificed for examination of the intestinal tract, and transit of chyme was determined as the distance from the distal duodenum edge to the leading edge of the carmine-stained area within the intestine. This distance was expressed as a percentage of the small intestine total length of an individual mouse, and amounted to 82.6±3.4% in the control group (n=10) and 62.7±3.6% in the experimental group (n=10) (P<0.001, Figure 7E). Three animals appeared to experience cessation of intestinal motility, which manifested as their refusal to eat despite the previous starvation. In this connection it is interesting to note that sensations of satiety and satiation inversely correlate with GI motility^28^.

## Discussion

We have characterised Pico145 as a novel GI-active small molecule that potently inhibits mI_CAT_ in single ileal myocytes (IC_50_ ∼3 pM), carbachol-induced [Ca^2+^]_i_ and contractile responses, as well as *ex vivo* (spontaneous and carbachol-induced) and *in vivo* postprandial small intestinal motility. In the latter tests, higher concentrations were needed to produce functional effects, possibly due to diffusional barriers and/or nonspecific binding of this lipophilic compound in tissues, consistent with the relatively low aqueous solubility and high plasma protein binding of Pico145^29^.

Under physiological conditions, activation of M2 and M3 receptors by the same agonist generates simultaneous, but different signalling events, which, acting concurrently, produce TRPC4 openings. Moreover, TRPC4 activity is also strongly voltage- and Ca^2+^-dependent. These properties make TRPC4 an important focal point, pivotal for cholinergic excitation-contraction coupling in the gut. It is thus plausible that Pico145 may complement the existing antimuscarinic agents for controlling hyperactivity of smooth muscles. Notably, cholinergic contractions of the detrusor similarly involve activation of TRPC4 channels^30^, while neuronal TRPC4 channels are involved in overactive bladder disease^31^. Thus, our results may have wider implications for smooth muscle pathophysiology.

We first characterised the inhibitory effect of Pico145 on mI_CAT_ in ileal myocytes. Several features of these effects are consistent with previous findings using HEK293 cells stably expressing tetracycline-regulated human TRPC4^24^. First, Pico145 demonstrated a much higher potency for carbachol- or GTPγS induced currents compared to EA-induced currents (Figure 5 *vs.* Figures 1-3). Indeed, the IC_50_ value of 3.1 pM for mI_CAT_ is comparable to 9-59 pM (depending on the membrane potential) for recombinantly expressed TRPC4 activated by sphingosine 1-phosphate acting via a G protein signalling pathway. In agreement with the electrophysiological data, carbachol-induced intracellular Ca^2+^ rises were inhibited with an IC_50_ value of 2.7 pM. This is in contrast to EA-induced currents, in which case 100 pM Pico145 inhibited the current by only 43% (Figure 5). These results are in line with Pico145 being a competitive antagonist or negative allosteric modulator of the direct TRPC4 agonist EA^24^. Second, the altered kinetics of mI_CAT_ deactivation at negative potentials (Figure 2D,E) suggest reduction of the channel mean open dwell time by Pico145. Third, voltage dependence of Pico145 action suggests that Pico145 binding site is located within the membrane potential field of TRPC4. This is consistent with recent cryo-EM data for structurally similar TRPC5 in complex with Pico145, showing that the interaction takes place between the transmembrane domains of two TRPC5 subunits within a lipid-binding site that is conserved within TRPC channels^27^. This mode-of-action is consistent with the lipophilic nature of Pico145 and with our observations that its inhibitory effect develops rather slowly and that its kinetics are highly variable in different cells. Potentiation of the inhibitory effect by membrane depolarisation may be therapeutically beneficial for the treatment of disorders of smooth muscle hyperactivity, if this is associated or caused by membrane depolarisation. The slow action of Pico145 could result from different factors: Pico145 may need to build up in the membrane and/or enter a site from inside the membrane. It also may need to displace a phospholipid in the binding site^27^. Several of these steps (movement of the phospholipid, entering of Pico145) may depend on channel state – therefore the kinetics could be different in different cell types, with different TRPC4 isoforms, and in different cellular conditions.

Pico145 (1 nM) strongly suppressed carbachol-induced contractile responses and spontaneous activity of longitudinal and circular smooth muscle layers. Similarly, Pico145 applied at 3 nM inhibited spontaneous and carbachol-initiated motility of an intact ileal segment. Our *in vivo* results (inhibition of intestinal transit by 24%) are quantitatively similar to the previous findings, where small intestinal transit quantified in TRPC4^-/-^/TRPC6^-/-^ mice showed reduction by ∼25% compared to wild-type mice^8^. Thus, the inhibitory effect of Pico145 on intestinal motility *in vivo* is as effective as *trpc4/trpc6* gene deficiency.

In conclusion, our results show that Pico145, at picomolar concentrations, inhibits TRPC4-mediated mI_CAT_ in mouse ileal cells activated in a physiologically relevant manner by carbachol or intracellular GTPγS, but less so when activated by EA. Pico145 is a high-quality small molecule inhibitor of TRPC4 channels in native cells, which is useful for further research regarding the biological functions of TRPC channels and, potentially, for the treatment of hyperactivity of visceral smooth muscles.

## Materials and Methods

### Animal Studies, Tissues and Cells Preparation

Animal studies were carried out in accordance with the recommendations of the European Convention for the Protection of Vertebrate Animals used for Experimental and other Scientific Purposes and approved by the Institutional Animal Care and Use Committees. The BALb/c two month-old male mice were humanely euthanized, and then the longitudinal layer of the ileum was dissected and cut into small pieces (1-2 mm). Single myocytes were isolated by mechanical trituration following enzymatic tissue treatment in a divalent-cation free solution containing collagenase type 1A, trypsin inhibitor II-S and bovine serum albumin (all at 1 mg/ml) at 36.8 °C for 18 min. Cell suspension was placed onto coverslips with addition of the normal modified Krebs solution at 2:1 ratio and stored in the fridge at 4 °C. Freshly isolated cells were used within 6-8 hours. All tissue and cell preparation procedures were carried out in a modified Krebs solution containing (in mM): 120 NaCl, 6 KCl, 2.5 CaCl_2_, 1.2 MgCl_2_, 12 D-glucose, 10 HEPES, pH adjusted to 7.35 with NaOH.

### Patch-clamp recordings

For whole-cell patch-clamp recordings, patch pipettes made of borosilicate glass (Sutter Instrument, Novato, CA, USA) had a resistance of 2.5-4 MΩ. The pipette solution contained (in mM): 80 CsCl, 1 MgATP, 4.6 CaCl_2_, 10 BAPTA, 5 creatine, 5 D-glucose, 10 HEPES, pH adjusted to 7.4 with CsOH. Intracellular Ca^2+^ concentration ([Ca^2+^]_i_) strongly buffered at 100 nM ensured mI_CAT_ stability, as this current is strongly Ca^2+^-dependent^16^. A gigaseal was formed in a normal modified Krebs solution, but before obtaining the whole-cell configuration for mI_CAT_ recordings, it was replaced by a Cs^+^ solution containing (in mM): 120 CsCl, 12 D-glucose, 10 HEPES, pH adjusted to 7.4 by CsOH. Membrane currents were recorded at room temperature (22-25 °C) using an Axopatch 200B voltage-clamp amplifier interfaced to a Digidata 1322A and a PC running the pClamp 8 program (Molecular Devices, San Jose, CA, USA). The data were filtered at 2 kHz and sampled at 10 kHz for storage and analysis. Our standard voltage protocol included a combination of voltage steps interposed with a slow voltage ramp from 80 to −120 mV, which were applied from the holding potential of −40 mV at 30 s intervals. This allowed simultaneous assessment of changes in mI_CAT_ kinetics (deactivation at −120 mV, activation at 80 mV, reactivation at −40 mV), as well as steady-state I-V relationships.

### Ca^2+^ imaging

Changes in [Ca^2+^]_i_ were measured using Fura-2. The dye was loaded by 30-min incubation of the cells with the cell-permeant Fura-2 AM (2 µM) followed by a 20-min wash-out period in a modified Krebs solution to allow time for de-esterification. Excitation wavelengths of 340 and 380 nm were used to monitor the fluorescence signals of the Ca^2^-bound and Ca^2+^-free dye. Fluorescence was recorded and analysed using an inverted Olympus IX51 microscope, Imaging station Cell^M^/Cell^R^ and the Olympus xCellence software (Olympus Optical Co., LTD, Nagano, Japan). [Ca^2+^]_i_ changes are reported as fluorescence ratio F_340_/F_380_ and cells were considered responsive if they demonstrated a change in fluorescence ratio >10% of baseline. All experiments were conducted at room temperature (∼22 °C).

### In vitro contraction recordings

Tensiometric recordings were carried out on both longitudinal and circular smooth muscle layers of the ileum. The longitudinal SM strips were obtained as described above. The circular muscle layer was dissected, washed, and cleaned of connective and adipose tissues and cut into 1 to 1.5-mm-wide rings. SM preparations were fixed between a stationary stainless steel hook and an isometric force transducer (AE 801, SensoNor A/S, Norten, Norway) coupled to an AD converter Lab-Trax 4/16 (World Precision Instruments, Inc., Sarasota, FL, USA). The data were continuously recorded using DataTrax2 (World Precision Instruments, Inc., Sarasota, FL, USA).

A flowing tissue bath maintained temperature at 37 °C and contained the modified Krebs-bicarbonate buffer (in mM): 133 NaCl, 4.7 KCl, 2.5 CaCl_2_, 1.2 MgCl_2_, 16.3 NaHCO_3_, 1.38 NaH_2_PO_4_, 10 HEPES, 7.8 D-glucose, pH adjusted to 7.4 with NaOH. Resting tension was adjusted to 0.5 g and 0.1 g for the longitudinal and circular muscles, respectively. Samples were allowed to equilibrate for 1 h under resting tension before the experiments commenced.

### Ex vivo motility test

The ileal preparation was cleaned from adipose tissues and placed in a normal modified Krebs solution maintained at 37 °C. Movements of intestinal segments were recorded using a digital camera. The contractile activity of the intact ileal segment was analyzed using the ImageJ software (National Institutes of Health, Bethesda, MA, USA) and the MTrackJ plug-in. Several points of interest were selected along the ileum in the first frame of the video, and then in every next frame the selected points were shifted according to the ileum movements, finally the tracks of points were created and processed by the software as shown in Supplementary Figure 1. Ileal contractions were expressed as the tracks of the selected points at one frame per 0.03 s. The measurements represent the D2S parameter, the distance (in pixels) from the start point of the track to the end point for each of the selected points.

### In vivo small intestinal transit

Evaluation of small intestinal transit was performed as described^8^ with some modifications. Mice were deprived of food for 24 h, but had free access to water supplemented with 20% glucose (w/v). 0.25 mL of distilled water with/without Pico145 was given by oral gavage using a flexible plastic needle. Pico145 aliquots were diluted in distilled water in such amount that its final dose was 1 mg per 1 kg body weight. The animals were fed 1 g of grains stained by carmine red (6% w/v) 10 minutes later and were sacrificed by cervical dislocation 30 min after feeding. The abdominal cavity was quickly dissected and the complete small intestine carefully removed and placed on white paper. Chyme displacement was observed as a carmine-stained area within the intestine, and intestinal transit was quantified as the distance from the pyloric sphincter to the forward edge of the carmine-coloured area. This distance was expressed as percentage of the small intestine total length of an individual mouse.

### Chemicals

(-)-Englerin A (EA) was obtained from PhytoLab GmbH & Co. KG (Vestenbergsgreuth, Germany). Pico145 was synthesized as described previously^24^. All other reagents, enzymes and salts were obtained from Sigma-Aldrich (St. Louis, MO, USA).

EA and Pico145 were dissolved in dimethyl sulfoxide (DMSO) to prepare their 10 mM stock solutions, aliquots of which were stored at −80 °C and −20 °C, respectively. Further dilutions to the required concentration were made with bathing solutions, or with distilled water for *in vivo* studies. As tested previously, DMSO at 1:800 v v^−1^ had negligible effect on mI_CAT_^11^, while at the maximal concentrations of EA or Pico145 used in the present study (i.e., 3 nM) DMSO dilution was at least 3.3:10^6^ v v^−1^; hence, no vehicle only controls were necessary.

### Data Analysis

Data were analyzed and plotted using the Clampfit 8 (Molecular Devices, Sunnyvale, CA, USA) and Origin 2022 software (OriginLab Corporation, Northampton, MA, USA). Data are given as means ± SEM, and *n* indicates the number of cells or smooth muscle samples tested. Differences between groups were evaluated using Student’s *t*-test after testing for their normal distribution. Differences were considered to be significant at P<0.05.

## Acknowledgments

This study was supported by the Ministry of Education and Science of Ukraine grant 0122U001535, National Academy of Science of Ukraine grant 0122U002126 for research laboratories/groups of young investigators in priority areas of Science and Technology for 2002-2023). Work by RSB at the University of Leeds was supported by the British Heart Foundation (PG/19/2/34084). The graphical abstract was created with BioRender.com.

## Abbreviations

EA: (-)-englerin A

GI: gastrointestinal

InsP_3_: D-myo-inositol 1,4,5-trisphosphate

mI_CAT_: muscarinic cation current

PIP_2_: phosphatidylinositol 4,5-bisphosphate

SM: smooth muscle

---

**Supplementary Figure 1.**
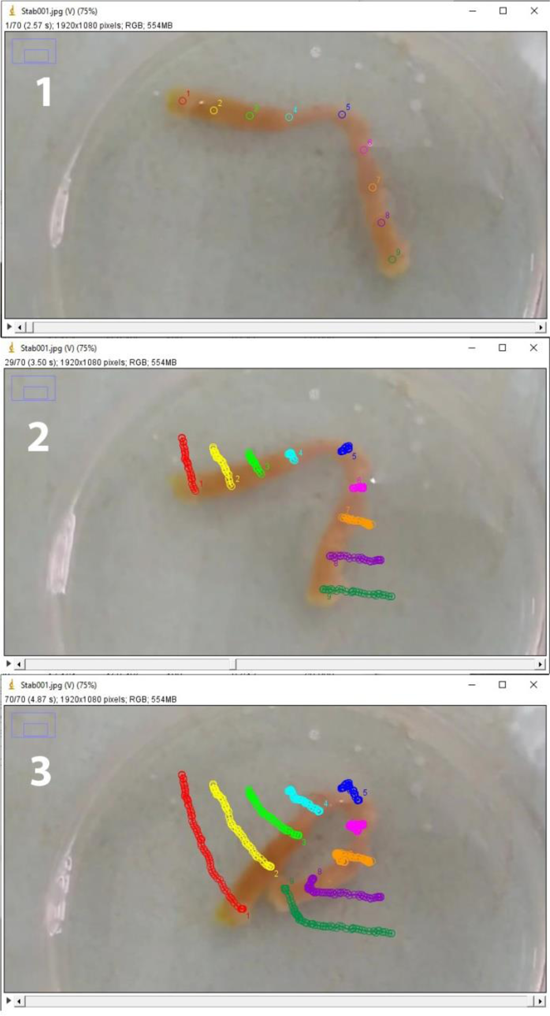
Analysis of contractile activity of an intact ileal segment (see Supplementary Video 1) using ImageJ and MTrackJ. 1 - the first frame of the analyzed video with 9 selected points along the preparation. 2 - randomly chosen frame of the analyzed video showing displacement of the 9 selected points. 3 - the last frame of the analyzed video showing the whole tracks of the 9 selected points.

## References

1. Bolton TB, Prestwich SA, Zholos AV, et al. Excitation-contraction coupling in gastrointestinal and other smooth muscles. Annu Rev Physiol 1999;61:85–115.

2. Sanders KM. Nerves, smooth muscle cells and interstitial cells in the GI tract: Molecular and cellular interactions. In: Rao SSC, Lee Y., Ghoshal UC, eds. Clinical and Basic Neurogastroenterology and Motility. Academic Press; 2020:3–16.

3. Zholos AV. Regulation of TRP-like muscarinic cation current in gastrointestinal smooth muscle with special reference to PLC/InsP_3_/Ca^2+^ system. Acta Pharmacol Sin 2006;27:833–842.

4. Unno T, Kwon SC, Okamoto H, et al. Receptor signaling mechanisms underlying muscarinic agonist-evoked contraction in guinea-pig ileal longitudinal smooth muscle. Br J Pharmacol 2003;139:337–350.

5. Kuriyama H, Kitamura K, Itoh T, et al. Physiological features of visceral smooth muscle cells, with special reference to receptors and ion channels. Physiol Rev 1998;78:811–920.

6. Beech DJ. Actions of neurotransmitters and other messengers on Ca^2+^ channels and K^+^ channels in smooth muscle cells. Pharmacol Ther 1997;73:91–119.

7. Zholos AV. Muscarinic effects on ion channels in smooth muscle cells. Neurophysiology 1999;31(3):173–187.

8. Tsvilovskyy VV, Zholos AV, Aberle T, et al. Deletion of TRPC4 and TRPC6 in mice impairs smooth muscle contraction and intestinal motility in vivo. Gastroenterology 2009;137(4):1415–1424.

9. Zholos AV, Bolton TB. Muscarinic receptor subtypes controlling the cationic current in guinea-pig ileal smooth muscle. Br J Pharmacol 1997;122(5):885–893.

10. Tanahashi Y, Katsurada T, Inasaki N, et al. Further characterization of the synergistic activation mechanism of cationic channels by M2 and M3 muscarinic receptors in mouse intestinal smooth muscle cells. Am J Physiol - Cell Physiol 2020;318:C514–C523.

11. Zholos A V., Tsytsyura YD, Gordienko D V., et al. Phospholipase C, but not InsP_3_ or DAG, - dependent activation of the muscarinic receptor-operated cation current in guinea-pig ileal smooth muscle cells. Br J Pharmacol 2004;141:23–36.

12. Yan H-D, Okamoto H, Unno T, et al. Effects of G-protein-specific antibodies and Gβγ subunits on the muscarinic receptor-operated cation current in guinea-pig ileal smooth muscle cells. Br J Pharmacol 2003;139(3):605–615.

13. Jeon JP, Hong C, Park EJ, et al. Selective Gαi subunits as novel direct activators of transient receptor potential canonical (TRPC)4 and TRPC5 channels. J Biol Chem 2012;287:17029– 17039.

14. Thakur DP, Tian J Bin, Jeon J, et al. Critical roles of Gi/o proteins and phospholipase C-δ1 in the activation of receptor-operated TRPC4 channels. Proc Natl Acad Sci U S A 2016;113:109– 1097.

15. Otsuguro KI, Tang J, Tang Y, et al. Isoform-specific inhibition of TRPC4 channel by phosphatidylinositol 4,5-bisphosphate. J Biol Chem 2008;283:10026–10036.

16. Gordienko DV, Zholos AV. Regulation of muscarinic cationic current in myocytes from guinea-pig ileum by intracellular Ca2+ release: A central role of inositol 1,4,5-trisphosphate receptors. Cell Calcium 2004;36(5):367–386.

17. Ehlert FJ, Pak KJ, Griffin MT. Muscarinic agonists and antagonists: effects on gastrointestinal function. Handb Exp Pharmacol 2012;208:343–374.

18. Kim YC, Kim SJ, Sim JH, et al. Protein kinase C mediates the desensitization of CCh-activated nonselective cationic current in guinea-pig gastric myocytes. Pflugers Arch - Eur J Physiol 1998;436:1–8.

19. Tsvilovskyy VV, Zholos AV, Bolton TB. Effects of polyamines on the muscarinic receptor-operated cation current in guinea-pig ileal smooth muscle myocytes. Br J Pharmacol 2004;143:968–975.

20. Bon RS, Wright DJ, Beech DJ, et al. Pharmacology of TRPC channels and its potential in cardiovascular and metabolic medicine. Annu Rev Pharmacol Toxicol 2022;62:427–446.

21. Wang H, Cheng X, Tian J, et al. TRPC channels: Structure, function, regulation and recent advances in small molecular probes. Pharmacol Ther 2020;209:107497.

22. Karuna Therapeutics announces exclusive global license agreement for Goldfinch Bio’s investigational TRPC4/5 product candidates. Business Wire 2023. Available at: https://www.businesswire.com/news/home/20230202005296/en/Karuna-Therapeutics-Announces-Exclusive-Global-License-Agreement-for-Goldfinch-Bio's-Investigational-TRPC45-Product-Candidates [Accessed July 5, 2023].

23. Walsh L, Reilly JF, Cornwall C, et al. Safety and efficacy of GFB-887, a TRPC5 channel inhibitor, in patients with focal segmental glomerulosclerosis, treatment-resistant minimal change disease, or diabetic nephropathy: TRACTION-2 Trial Design. Kidney Int Reports 2021;6:2575–2584.

24. Rubaiy HN, Ludlow MJ, Henrot M, et al. Picomolar, selective, and subtype-specific small-molecule inhibition of TRPC1/4/5 channels. J Biol Chem 2017;292:8158–8173.

25. Just S, Chenard BL, Ceci A, et al. Treatment with HC-070, a potent inhibitor of TRPC4 and TRPC5, leads to anxiolytic and antidepressant effects in mice. PLoS One 2018; 13(1):e0191225.

26. Zholos AV, Zholos AA, Bolton TB. G-protein-gated TRP-like cationic channel activated by muscarinic receptors: Effect of potential on single-channel gating. J Gen Physiol 2004;123(5):581–598.

27. Wright DJ, Simmons KJ, Johnson RM, et al. Human TRPC5 structures reveal interaction of a xanthine-based TRPC1/4/5 inhibitor with a conserved lipid binding site. Commun Biol 2020;3:1–11.

28. Janssen P, Berghe P Vanden, Verschueren S, et al. Review article: the role of gastric motility in the control of food intake. Aliment Pharmacol Ther 2011;33:880–894.

29. Yu M, Ledeboer MW, Daniels M, et al. Discovery of a potent and selective TRPC5 inhibitor, efficacious in a focal segmental glomerulosclerosis model. ACS Med Chem Lett 2019;10:1579–1585.

30. Griffin CS, Bradley E, Dudem S, et al. Muscarinic receptor induced contractions of the detrusor are mediated by activation of TRPC4 channels. J Urol 2016;196:1796–1808.

31. Boudes M, Uvin P, Pinto S, et al. Crucial role of TRPC1 and TRPC4 in cystitis-induced neuronal sprouting and bladder overactivity. PLoS One 2013; 8(7):e69550.

